# Borna disease virus 1 infection causing fatal meningoencephalomyelitis in wild European hedgehogs in known endemic areas, Germany, 2022 to 2024

**DOI:** 10.1101/2025.07.08.663648

**Authors:** Effrosyni Michelakaki, Benjamin Schade, Brigitte Boehm, Eva Kappe, Marcel Suchowski, Anne Kupca, Magdalena Schumacher, Anna Maria Gager, Friederike Liesche-Starnecker, Zoltan Bago, Andreas Blutke, Martin Beer, Dennis Rubbenstroth, Kaspar Matiasek

**Affiliations:** Institute of Veterinary Pathology, Centre for Clinical Veterinary Medicine, Ludwig Maximilians-Universität München, Munich, Germany; Bavarian Animal Health Service, Poing, Germany; Bavarian Health and Food Safety Authority, Oberschleißheim, Germany; Department of Neuropathology, Pathology, Medical Faculty, University of Augsburg, Augsburg, Germany; AGES, Institute for Veterinary Disease Control, Mödling, Austria; Institute of Diagnostic Virology, Friedrich-Loeffler-Institut, Greifswald-Insel Riems, Germany

**Keywords:** encephalitis, emerging, Erinaceus europaeus, zoonosis, insectivore, Eulipotyphla, host, BoDV-1, lethal, endemic

## Abstract

Herein, we report Borna disease virus 1 (BoDV-1) infection in seven wild European hedgehogs *(Erinaceus europaeus)* from an endemic area in Germany. BoDV-1 causes encephalitis with a fatality rate of more than 90% in domestic mammals and humans.

Currently, the bicolored white-toothed shrew (*Crocidura leucodon*) is the only known reservoir host. As hedgehogs are distant relatives of shrews and often cared for by humans, the cases raise concern regarding a potential zoonotic risk.

All of the hedgehogs that tested positive for BoDV-1 had succumbed to neurological disease and exhibited severe polio-predominant lymphoplasmohistiocytic meningoencephalitis.

However, due to the detection of viral antigens in non-neural cells in one animal, it cannot be completely excluded that some infected hedgehogs shed the virus. Although direct BoDV-1 transmission is known to be inefficient, our results emphasize the necessity of hygiene measures when handling hedgehogs, especially those with neurological signs from BoDV-1-endemic regions.

## Introduction

Borna disease virus 1 (BoDV-1; species *Orthobornavirus bornaense*, family *Bornaviridae*) is known for decades as the causative agent of Borna disease, a usually fatal, immune-mediated meningoencephalitis continuously identified throughout endemic areas covering parts of Germany, Austria, Liechtenstein and Switzerland (*1*). Borna disease has been demonstrated to affect a broad range of domestic mammals, particularly horses, alpacas and sheep (*1–4*). Since 2018, it has also been proven that BoDV-1 can cause encephalitis in humans. Up to six cases are reported each year, with a case fatality rate of over 90% (*1, 5–7*). Domestic mammals and humans are known to serve as dead-end hosts, in which the virus possesses an almost exclusively neurotropic tissue distribution without detectable viral shedding (*3, 8*). Their infection was demonstrated to result from spill-over transmission from a natural reservoir (*1, 9, 10*). Currently, the insectivorous bicolored white-toothed shrew (*Crocidura leucodon*) is the only known reservoir host species (*11–13*). Infected shrews develop life-long viral persistence with a broad tissue distribution and continuous viral shedding, but no apparent tissue lesions or clinical disease (*11*).

The European hedgehog *(Erinaceus europaeus)* is another insectivorous species indigenous to Europe that often comes into close contact with humans, particularly when being cared for during hibernation in private households and rescue centers. Although hedgehogs have been associated with various zoonotic diseases (*14, 15*), there have been no reports on BoDV-1 infection so far.

Herein, we present the findings relating to BoDV-1 infection and fatal encephalitis in seven wild European hedgehogs from an endemic area in Germany. The first case was detected in 2022. BoDV-1 was identified by reverse transcriptase quantitative polymerase chain reaction (RT-qPCR) in the brains of these animals. Previous testing for various encephalitic pathogens reported in this or other species, such as tick-borne encephalitis virus (TBEV) (*16*), rabies virus (*17*), canine distemper virus (CDV) (*18*) and rustrela virus (RusV) (*19*), had yielded negative results. After the diagnosis of the second case in May 2024 the awareness amongst hedgehog rescue centers and diagnostic institutions in the region increased considerably and led to identification of a series of additional cases.

The taxonomic proximity of hedgehogs with shrews (*20*), the known BoDV-1 reservoir hosts, raised concerns about hedgehogs being able to shed the virus and thereby lead to potential human exposure. Therefore, a comprehensive analysis of the confirmed cases was performed. This included detailed histopathology and extensive characterization of the viral tissue distribution and cell tropism using immunohistochemistry (IHC) and RNAscope® *in situ* hybridization (ISH) as well as phylogeographic analysis of hedgehog-derived BoDV-1 sequences. Additionally, a screening of non-encephalitic hedgehogs from endemic areas was initiated and is still ongoing.

## Methods

### Case selection, sample collection and diagnostic investigations

This investigation focused on eight wild hedgehogs that died or were euthanized due to a devastating course of encephalitis, histopathologically diagnosed on postmortem examination as described below. In addition, 23 deceased non-encephalitic hedgehogs from endemic regions in Bavaria were included in the study serving as controls (Supplemental table S1).

All animals underwent a complete postmortem examination, and a broad set of organs and tissues was fixed in 10% neutral buffered formalin for a minimum of 24 hours for histopathological analysis. From most cases, brain tissue and additional samples were snap-frozen for further investigation. Additionally, an oral swab and a fecal sample were collected from case 7.

Fresh-frozen or formalin-fixed paraffin-embedded (FFPE) brain tissue of all animals were tested using a BoDV-1-specific RT-qPCR for BoDV-1 RNA (*1, 3*). Furthermore, all available fresh-frozen samples and swabs of BoDV-1-positive animals were tested.

For all BoDV-1-positive animals, staining for BoDV-1 antigen and RNA was performed by IHC and RNAscope® ISH as described below.

Brain samples from encephalitic animals were additionally tested for other known encephalitic viruses, including TBEV and RusV by RT-qPCR (Supplemental table S2) (*21, 22*) and rabies virus (*23*) and CDV (*24*) by IHC as described in the appendix, section B. **Screening for BoDV-1 antigen distribution via immunohistochemistry**

IHC for BoDV-1 nucleoprotein (N) was performed using mouse monoclonal antibody Bo18 (*25*) on all sections of all available tissues. Rabbit anti-BoDV-1 N polyclonal hyperimmune serum #201 (*3*) was also used on selected central nervous system (CNS) and peripheral non-neural tissue sections of case 5. All procedures including detailed information on employed antibodies can be retrieved from the appendix (Appendix, section B). A BoDV-1-positive horse brain was used as positive control. Distinctive multiple and coalescing DAB-brown intracellular dots, as seen in the positive control, were considered positive.

### Screening for BoDV-1 RNA tissue distribution via RNAscope® in situ hybridization

Tissue sections of the CNS of all BoDV-1-positive animals, as well as a selection of peripheral organs underwent RNAscope® ISH for BoDV-1 RNA (V-BoDV1-G targeting viral RNA encoding for the matrix protein (M) and glycoprotein (G) genes; genome positions 2,236 to 3,747of BoDV-1 reference genome NC_001607.1; RNAscope®, Advanced Cell Diagnostics, Inc., USA) as described previously (*26*).

### Lesion characterization

Formalin-fixed tissues were macroscopically evaluated before and after fixation /trimming. The brain was trimmed at multiple planes. Representative areas of telencephalon, diencephalon, brain stem and cerebellum were processed for microscopic examination. Spinal cord sections comprised transverse and longitudinal sections upon decalcification of the vertebral column by a 20% EDTA solution. Representative sections were taken from all available, adequately preserved peripheral organs and tissues. All samples underwent an ascending alcohol series up to xylene using an automatic histoprocessor. Thereafter, the samples were embedded in paraffin, cut into 2-4 μm-thick sections and all sections were stained with hematoxylin and eosin (H&E) stain for routine microscopic examination (*27*).

To phenotype the inflammatory cell infiltrations, IHC was carried out by staining for T lymphocyte marker CD3, B lymphocyte marker Pax 5, and macrophage/microglial marker Iba1. The degree and distribution of gliosis was highlighted using the astrocyte marker glial fibrillary acidic protein (GFAP). All procedures including detailed information on employed antibodies can be retrieved from the appendix, section B.

### RNA extraction and RT-qPCR testing for BoDV-1

RNA from fresh-frozen tissue samples and swabs was extracted by using the NucleoMag VET kit (Macherey-Nagel, Düren, Germany) with a KingFisher Flex Purification System (Thermo Fisher Scientific, Bremen, Germany). The RNeasy FFPE kit (Qiagen, Hilden, Germany) was used for extraction of RNA from FFPE tissue. Procedures were performed as previously described (*1*). Semiquantitative detection of BoDV-1 RNA was performed by RT-qPCR assays BoDV-1 Mix-1 and Mix-6 (Supplemental table S2), as described previously (*1, 7*). RT-qPCR results are presented as cycle of quantification (Cq) values. An RNA preparation of BoDV-2 isolate No/98 (GenBank accession number AJ311524.1) was used as positive control and for calibration of the Cq values.

For all BoDV-1-positive animals, determination of partial BoDV-1 genome sequences covering at least the N, accessory protein (X) and phosphoprotein (P) genes (1,824 bases, positions 54 to 1,877 of reference genome U04608.1) were determined by Sanger sequencing of overlapping conventional RT-PCR products as described previously (*1, 3*). BoDV-1 sequences generated in this study were deposited in GenBank under accession numbers (PV357162.1 to PV357168.1). Phylogenetic analysis was performed using Geneious Prime 2021.0.1 (Biomatters Ltd., Auckland, New Zealand). A Neighbor Joining tree was calculated using the Jukes-Cantor model of all seven hedgehog-derived BoDV-1 sequences together with 258 N-X/P sequences from naturally infected animals and humans available from public databases (*1, 9*). The sequence of isolate BoDV-2 No/98 (AJ311524.1) was used to root the tree.

## Results

### Diagnostic testing, time of appearance, geographic origin & clinical presentation

Eight of the 31 evaluated hedgehogs from Bavaria showed lymphoplasmohistiocytic meningoencephalitis and BoDV-1 RNA was detected in the brains of seven of them (Table), but not in any of the 23 non-encephalitic hedgehogs (Supplemental table S1). Differential diagnostic testing by IHC and/or RT-qPCR did not detect TBEV, RusV, CDV or rabies virus in any of the encephalitis cases (data not shown).

**Table 1.** Overview of BoDV-1-infected European hedgehogs included in this study.

|  | Case 1 | Case 2 | Case 3 | Case 4 | Case 5 | Case 6 | Case 7 |
| --- | --- | --- | --- | --- | --- | --- | --- |
| District | Rottal-Inn | Ebersberg | Rottal-Inn | Ebersberg | Rottal-Inn | Ebersberg | Rottal-Inn |
| Rescue centre | A | Private person | A | B | Private person | B | A |
| Day of being found | 22/06/22 | 30/04/24 | 01/07/24 | 22/06/24 | 23/06/24 | 18/06/24 | 11/09/24 |
| Day of death | 16/07/22 | 05/05/24 | 10/07/24 | 12/07/24 | 17/07/24 | 15/08/24 | 12/09/24 |
| Survival time from admission (in days) | 24 | 6 | 9 | 20 | 24 | 58 | 1 |

The first BoDV-1-infected animal had died in July 2022, while the remaining six BoDV-1-positive cases occurred from May to September of 2024 (Table). All BoDV-1-positive hedgehogs originated from two different administrative districts within the BoDV-1-endemic area in Bavaria (Rottal-Inn and Ebersberg) and they were submitted for necropsy by two different hedgehog rescue centers or by private persons (Table). At necropsy, the hedgehogs weighed between 480g and 960g. Two were female and five were male. All of them had already exhibited neurological signs by the time they were found. According to the information provided by the submitters, all of the animals presented with incoordination and/or gait abnormalities. Seizures, spontaneous muscle twitching and vestibular signs with unilateral head tilt were observed in one or two animals, each. Three hedgehogs became apathic, two of which presented with impaired thermoregulation (Supplemental table S3). As the clinical signs progressed and the animals were not responsive to the administered treatments, all hedgehogs were euthanized due to poor prognosis or died after one to 58 days (Table).

### BoDV-1-associated lesion patterns

Apart from euthanasia-related changes, no macroscopical alterations were seen, including the central or peripheral nervous system (PNS) at necropsy of BoDV-1-infected hedgehogs.

Histologically, all seven BoDV-1-positive animals presented with a generalized angiocentric lymphoplasmohistiocytic meningoencephalitis (n=2) or meningoencephalomyelitis (n=5). The inflammatory infiltrates were widespread throughout all CNS regions of all seven animals (Figure 1) and multifocally invading the subarachnoid space, choroid plexus stromata and neuroparenchyma (Figures 2A-D). Also, microglial activation and astrogliosis (highlighted by Iba-1 or GFAP staining, respectively) were most extensive in the grey matter of cerebral cortices, diencephalon and brainstem (Figures 2B and 2F). All cases further featured a mild to moderate intralesional edema (Figure 2A).

**Figure 1.**
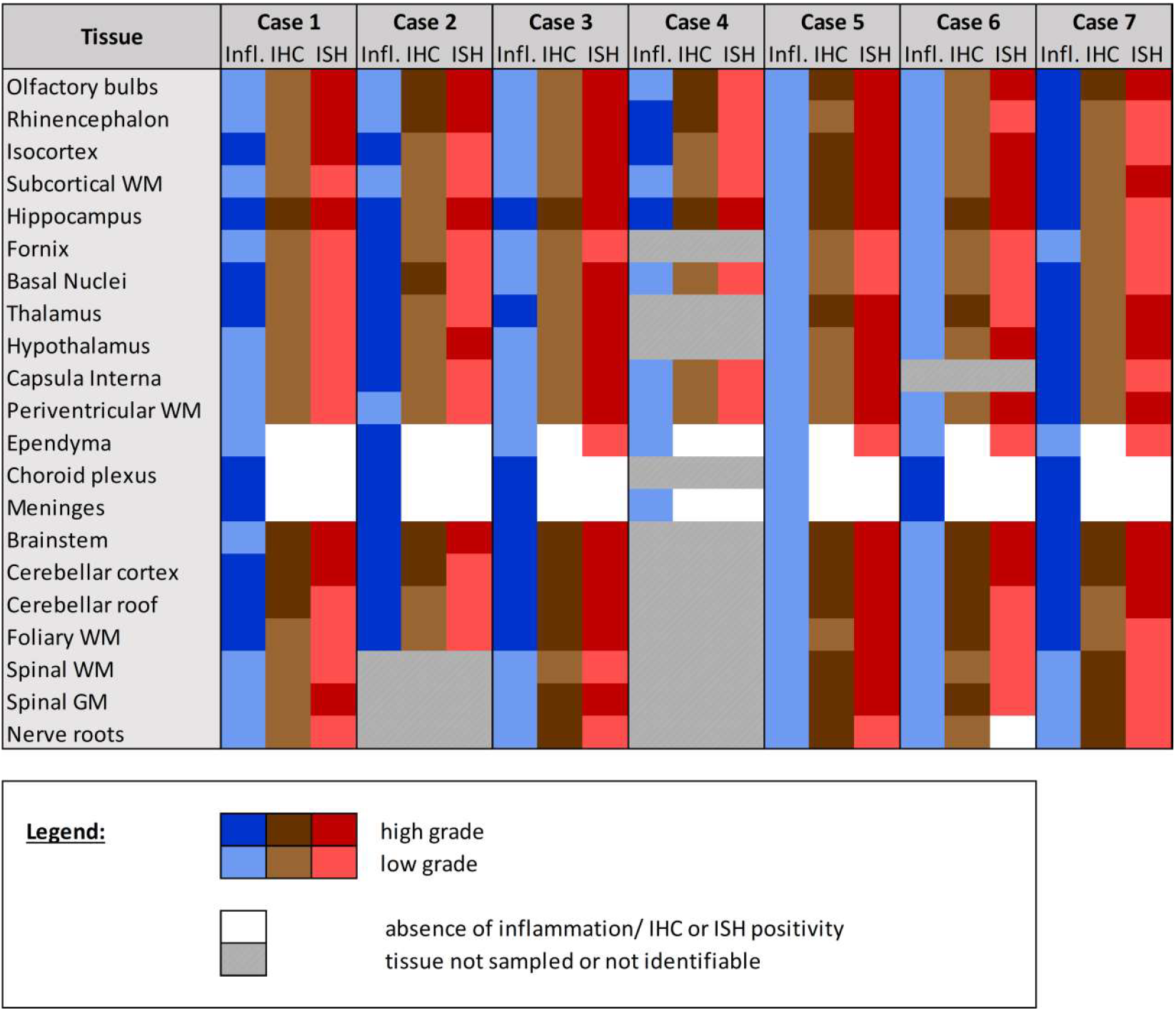
Inflammatory lesions and BoDV-1 antigen and RNA detection in the CNS of BoDV-1-infected European hedgehogs. Abbreviations: Infl. = inflammation; IHC = immunohistochemistry; ISH = RNAscope® ISH; WM = white matter; GM = grey matter.

**Figure 2.**
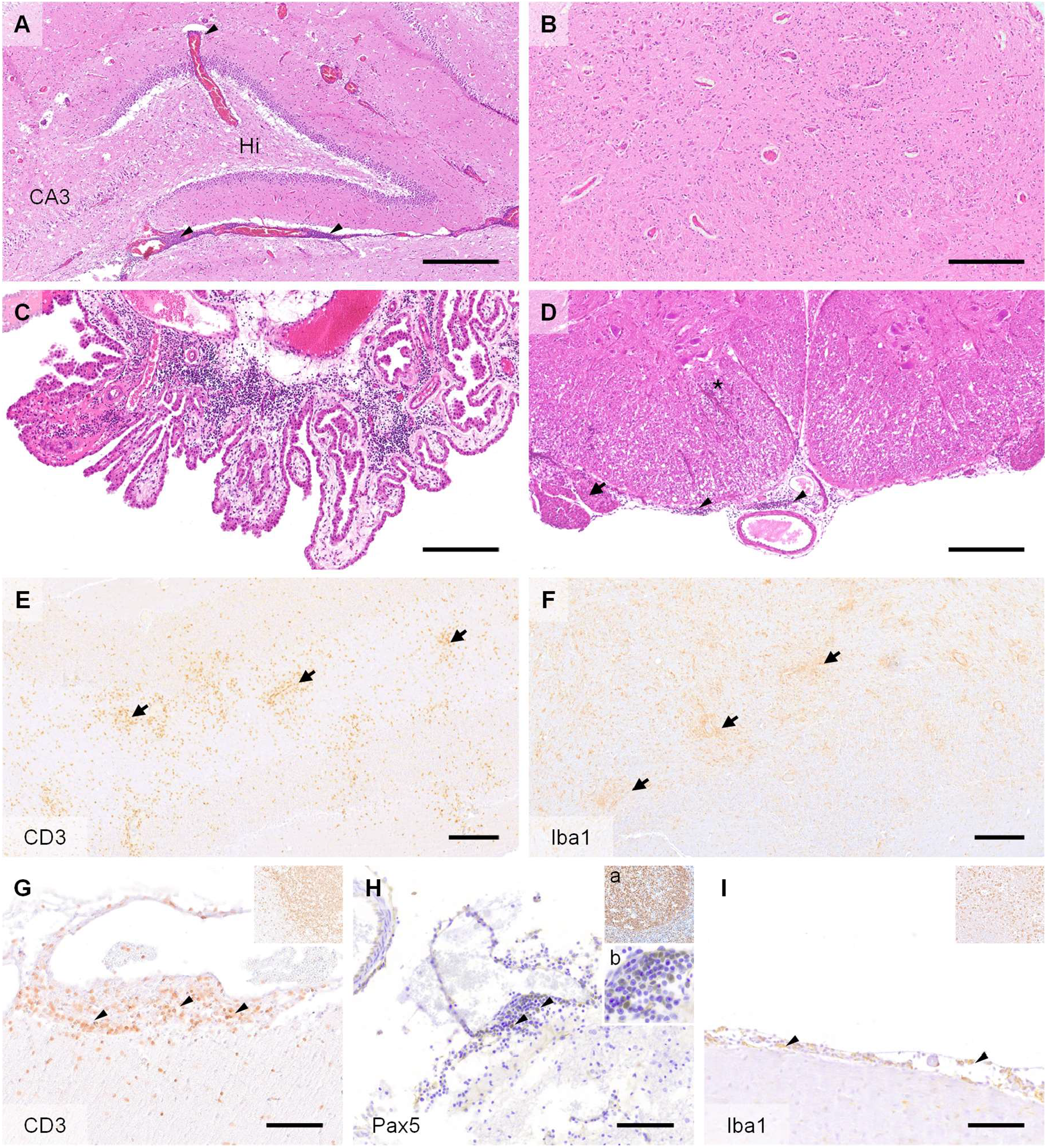
Representative changes and composition of immune cells in the CNS of BoDV-1-infected hedgehogs. A) Multifocal lymphohistiocytic infiltrates with perivascular cuff formation (arrowheads) and oedema extending from the hilus of dentate gyrus (Hi) towards CA3 segment of hippocampus proper (CA3). B) Typical scattered lymphocytic infiltrates intermingling with astro- and microgliosis within the brainstem. C) Infiltrates also extend into the choroid plexus. D) the subarachnoid space (arrowheads), nerve roots (arrow) and spinal white matter (asterisk). E) CD3-positive T lymphocytes (arrows) comprise the majority of invading immune cells throughout parenchyma and (G) meninges (arrowheads). F) The second most prominent cells involved in antiviral responses are Iba1-positive microglial cells and macrophages (arrows). H) B lymphocytes (arrowheads) resemble a minority of immune cells that mostly are concentrated around blood vessels and within the subarachnoid spaces. I) Subarachnoid spaces also contain multiple Iba1-positive macrophages (arrowheads). Insets in figures G and I, as well as inset Ha show the positive controls for each respective immunohistochemical staining (magnification ×20). Inset Hb shows a magnification of the positive B lymphocytes in figure H (magnification ×40). Origin of sections: A: Case 2. B: Case 7. C: Case 6. D: Case 5. E: Case 7. F: Case 7. G: Case 7. H: Case 1. I: Case 7. Scale bars: A: Bar=500 μm. B: Bar= 250 μm. C: Bar= 250 μm. D. Bar= 250 μm. E: Bar=200 μm. F: Bar=200 μm. G: Bar=100 μm. H: Bar=100 μm. I: Bar=100 μm. Stain: A-D: H&E. E-I: DAB, haematoxylin counterstain, IHC using markers: E: CD3, F: Iba1, G: CD3, H: Pax5, I: Iba1.

IHC-based phenotyping of inflammatory cells showed the affected brains to be extensively infiltrated predominantly by CD3-positive T lymphocytes, occasionally forming perivascular cuffs of up to 3–4-layer thickness (Figures 2E and 2G). Inflamed zones also showed moderate numbers of Iba1-positive macrophages and activated Iba1-positive microglial cells (Figures 2F and 2I) as well as GFAP-positive astrocytes, but only few scattered and mostly perivascular Pax5-positive B cells (Figure 2H). In cases 5 and 7, mild intraaxial vasculitic features were seen in addition to the infiltration of the neuroparenchyma.

The spinal cord was similarly, but overall, less severely affected, featuring multifocal infiltration of similar cellular composition mostly within the subarachnoid space (Figure 2D). Notably, a mild to moderate infiltration and mild to moderate multifocal spongiosis of spinal white matter was seen in all animals, even in areas with spared grey matter (Figure 2D). The inflammatory infiltrates also extended into the adjacent nerve roots and dorsal root ganglia (Figures 1 and 2D). Large fascicular nerves, distal and intramural ganglia and nerve branches showed minimal to no inflammation, except for a mild to moderate, focally extensive, lymphocytic infiltration of the cranial mesenteric ganglia and nerves in case 3.

Intranuclear Joest-Degen inclusion bodies were not observed.

Concurrent pathologies observed in the BoDV-1-positive hedgehogs are summarized in the appendix section C.

### Cell tropism and tissue distribution of BoDV-1 RNA and antigen

Moderate to high levels of BoDV-1 RNA (Cq values 16.6 to 24.8) were detected in the brains of all seven BoDV-1-positive animals and also in the spinal cord when available (Figure 3). Variable sets of fresh-frozen peripheral organs were available from five animals.

**Figure 3.**
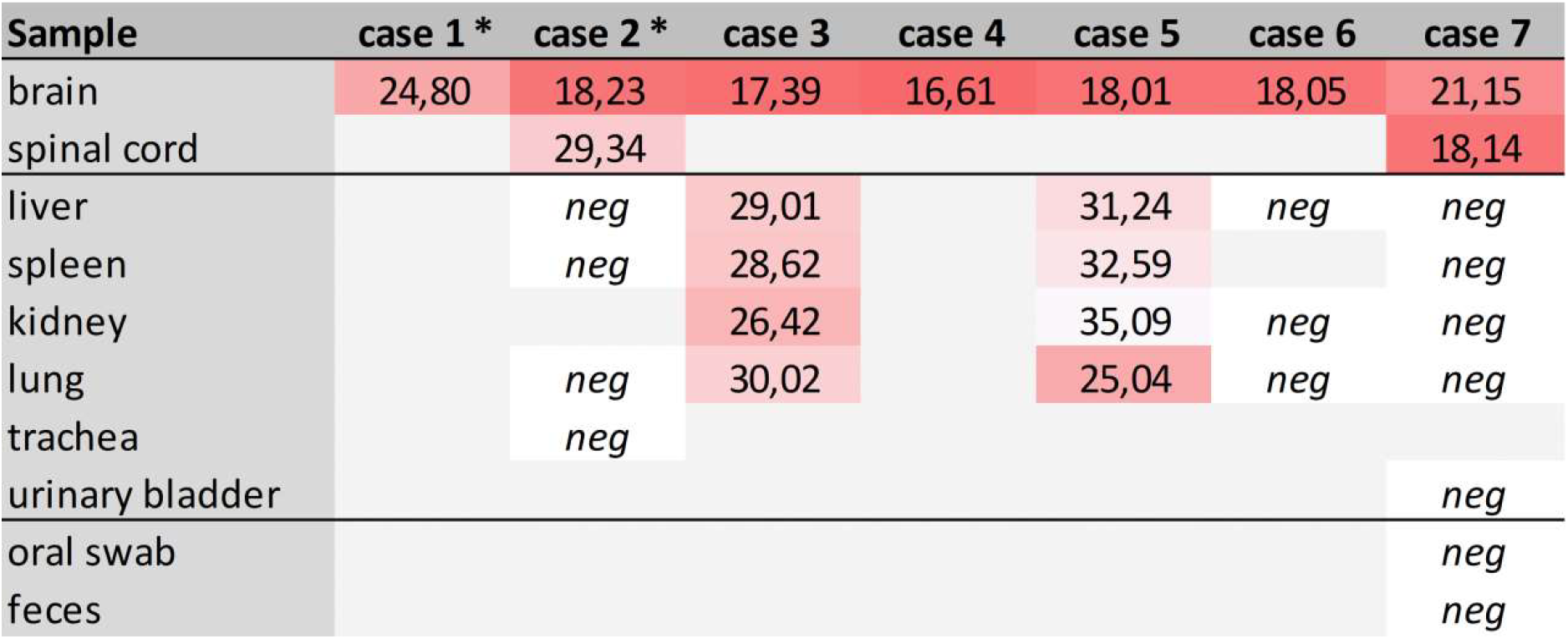
Detection of BoDV-1 RNA by RT-qPCR in available fresh-frozen neural and extra-neural tissues and additional samples collected postmortem. RT-qPCR results are presented as Cq values. *) Only formalin-fixed paraffin-embedded (FFPE) brain tissue was available for cases 1 and 2.

While no viral RNA was detectable in three of them, cases 3 and 5 had low to moderate BoDV-1 RNA levels detectable in all four tested tissue samples (liver, spleen, kidney and lung), with kidney or lung being most strongly positive (Cq 26.4 or 25.0, respectively; Figure 3). Postmortem oral swabs and fecal samples were collected only for case 7, which did not have detectable viral RNA in its periphery. Both samples were negative (Figure 3).

IHC using the Bo18 antibody and RNAscope® ISH revealed widespread cytoplasmic and nuclear positivity for the BoDV-1 N protein and genomic RNA, respectively, across neurons and glial cells without consistent hot spots in the brains and spinal cords of all BoDV-1-infected animals (Figures 1 and 4). Almost diffuse reactivity was seen throughout the neuroparenchyma, including white matter areas. IHC and ISH results were in agreement with each other, with the exception of the ependymal layer, in which only RNA was detected in four animals (Figure 4G). Neither antigen nor RNA were detected in the choroid plexus and meninges (Figure 1).

**Figure 4.**
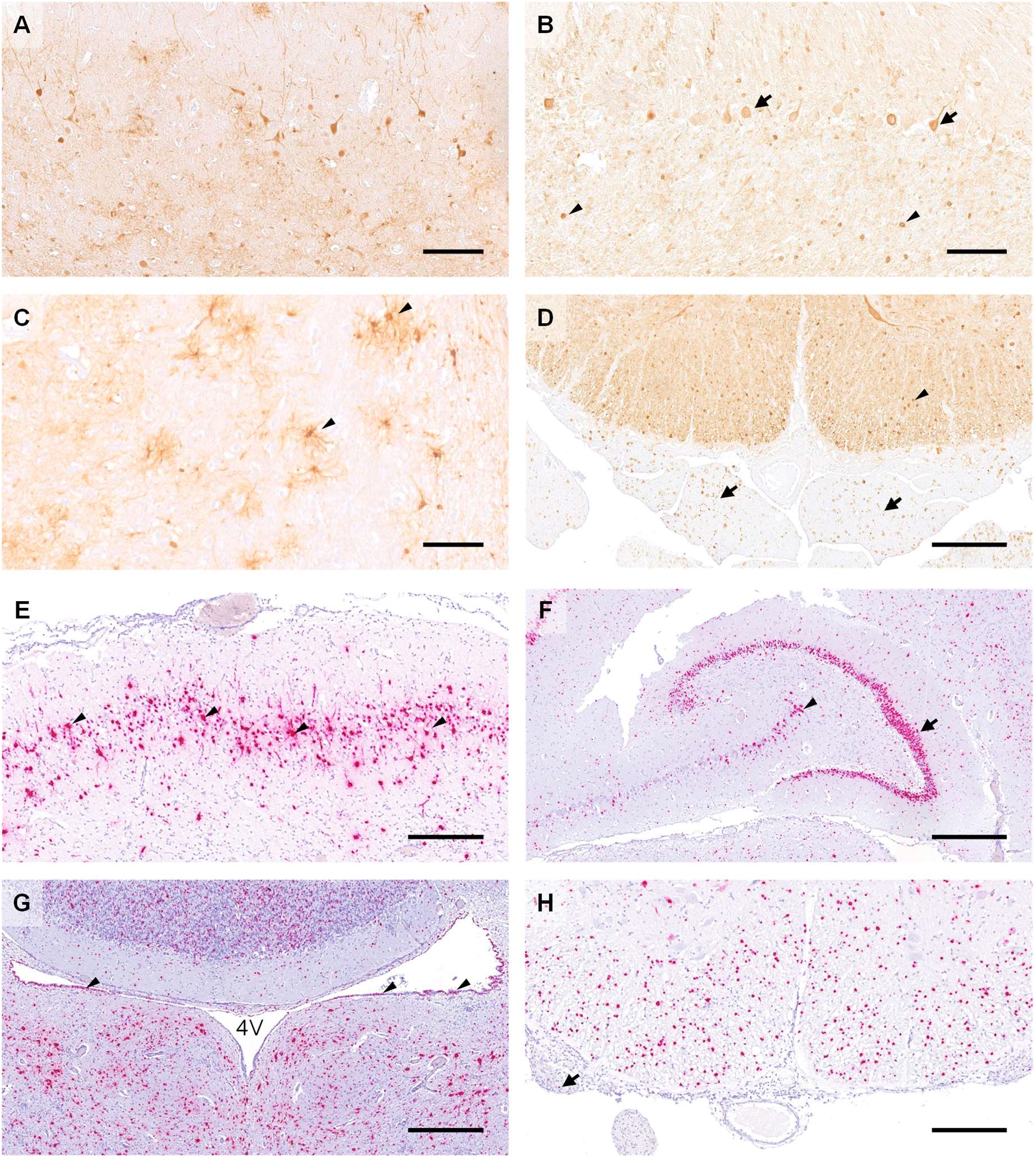
Distribution of virus antigen detected by immunohistochemistry using Bo18 antibody (A-D) and virus RNA detected by RNAscope® ISH (E-H) within the CNS of BoDV-1-infected hedgehogs. A) Multiple immunopositive neurons within the cerebral cortex. G) Multiple immunopositive Purkinje (arrows) and few granule cells (arrowheads). H) Multiple positive astroglial cells within the spinal white matter (arrowheads). I) Multiple positive oligondendroglial cells within the spinal white matter (arrowhead), as well as multiple positive nerve fibers within the adjacent nerve roots (arrows). E) Numerous ISH-positive neurons within the cortical layers of rhinencephalon (arrowheads). F) Granule cells of dentate gyrus (arrow) are almost entirely ISH-positive as do multiple neurons of cornu ammonis (arrowhead). G) Numerous ISH-positive neurons, glial as well as ependymal cells (arrowheads) surrounding the fourth ventricle (4V). H) Numerous ISH-positive oligodendroglial cells within the spinal white matter and rare signals within nerve roots (arrow). Origin of sections. A. Case 5. B: Case 3. C: Case 6. D: Case 5. E: Case 2. F: Case 5. G. Case 7. H. Case 5. Scale bars: A: Bar=100 μm. B. Bar=100 μm. C. Bar=100 μm. D. Bar=250 μm. E: Bar=250 μm. F: Bar=500 μm. G. Bar=500 μm. H. Bar=250 μm. Stain, A-I: DAB, hematoxylin counterstain. E-H: 2.5 HD Assay – RED, hematoxylin counterstain.

In addition, peripheral nerves and organs were analyzed by IHC. Positive signals were obtained from fascicular nerve roots (both dorsal and ventral), the dorsal root ganglia, peripheral nerves, distal and intramural ganglia (Figures 4D and 6A-H). Antigen-positive peripheral nerve branches and ganglia were seen within various organs, such as lungs (n=6), kidneys (n=4), mediastinum, spleen (both n=3), cranial mesenterium, intestine, the paravertebral musculature (all n=2) and salivary glands (n=1) (Figures 5 and 6A-H). Viral antigen was detected in ganglion and satellite cells, axons, myelin sheath and Schwann cells of the PNS. Additionally, the chromaffin cells of the adrenal medulla in cases 3 and 5 were diffusely and strongly positive (Figures 5 and 7A). Unlike its widespread presence in the PNS, specific detection of viral antigen in non-neural tissues was only seen in case 5, which exhibited a strong cytoplasmic immunopositivity in a single focus of renal tubular epithelial cells (Figures 5 and 6I).

**Figure 5.**
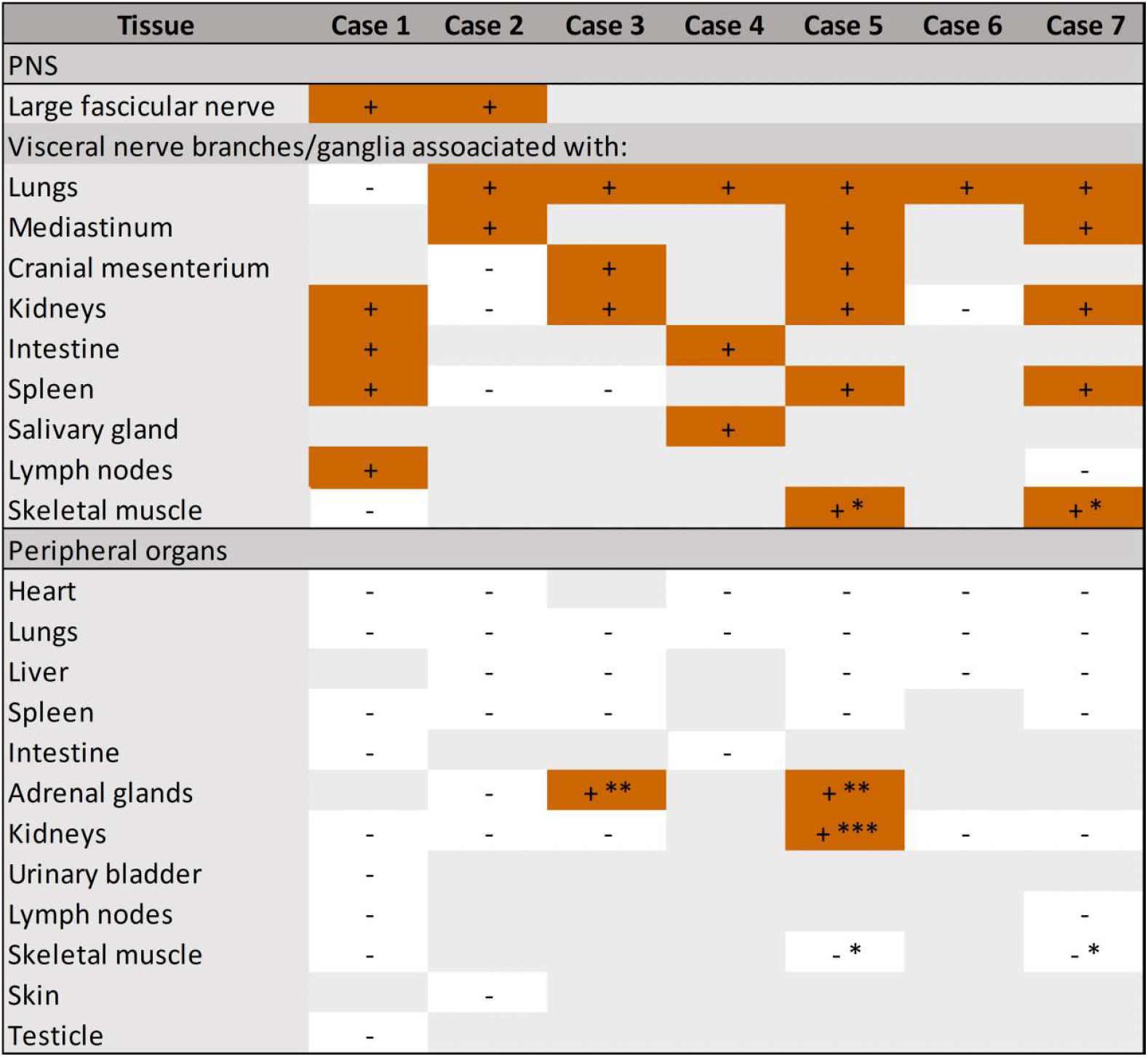
BoDV-1 antigen detection by immunohistochemistry using Bo 18 in peripheral nerves and peripheral organs of European hedgehogs. *= paravertebral musculature, **= chromaffin cells, ***= one focus of renal tubular epithelial cells. Nerve branches of organs that were not positive in any animal are not listed.

**Figure 6.**
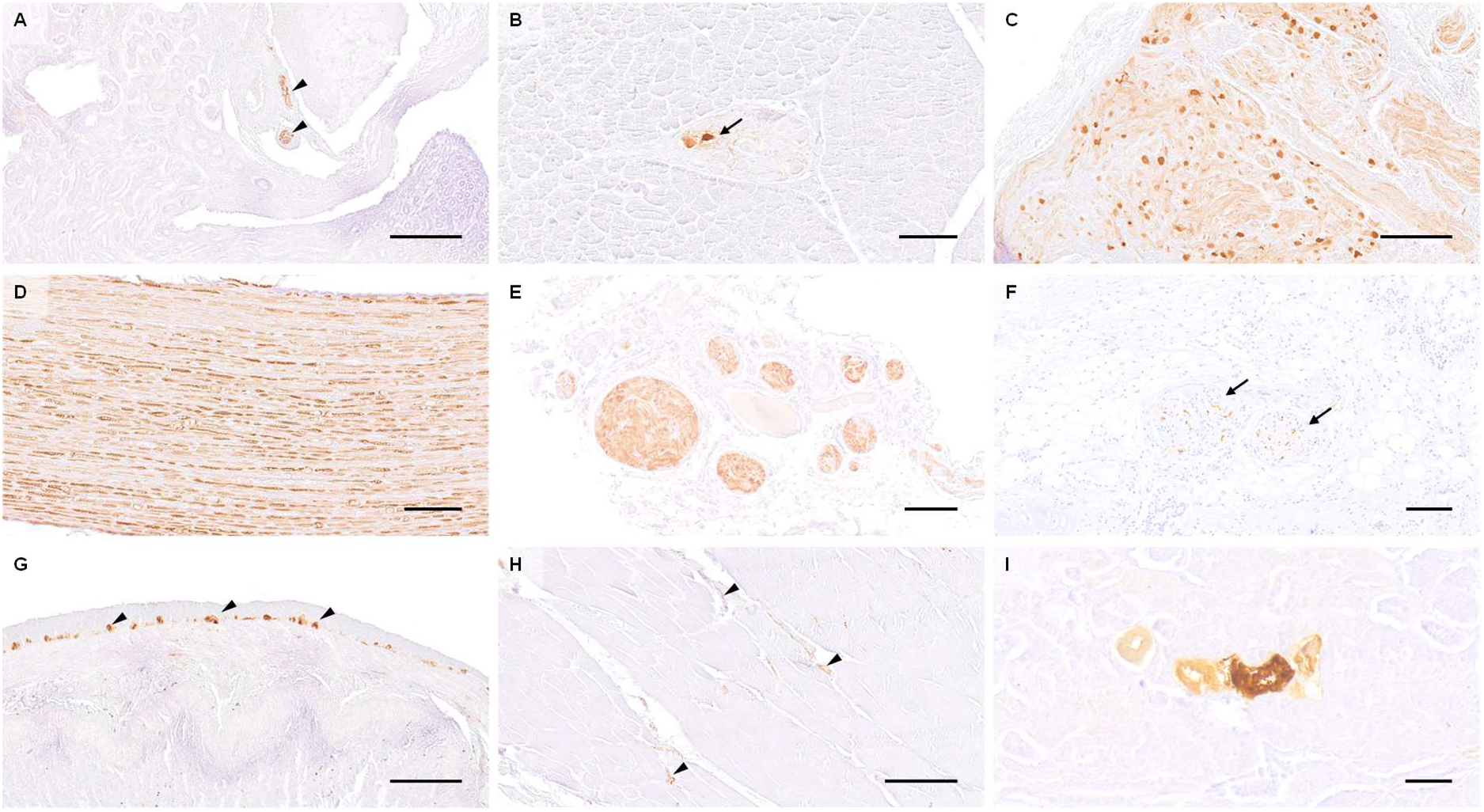
Distribution of virus antigen detected by immunohistochemistry using Bo18 antibody among peripheral organs and tissues of BoDV-1-infected hedgehogs. A) Immunopositive nerve branches within the renal pelvis (arrowheads). B) Immunopositive ganglion neurons within salivary gland (arrow). C) Marked positivity within cranial mesenteric ganglion. D) Diffusely immunopositive nerve fibers within a large fascicular nerve. E) Large immunopositive visceral nerve branches within the mediastinum. F) Pulmonary nerve branches with positive fibers (arrows). G) Almost diffusely immunopositive myenteric plexus (arrowheads). H) Immunopositive intramuscular nerve branches within the paravertebral musculature (arrowheads). I) A small group of immunopositive renal tubules. Origin of sections: A. Case 3. B. Case 4. C: Case 3. D. Case 1. E. Case 2. F. Case 7. G. Case 4. H: Case 5. I: Case 5. Scale bars: A. Bar=250 μm. B: Bar=100 μm. C: Bar=250 μm. D: Bar=100 μm. E. Bar=200 μm. F. Bar=100 μm. G. Bar=250 μm. H: Bar=250 μm. I: Bar=50 μm. Stain, A-I: DAB, hematoxylin counterstain.

**Figure 7.**
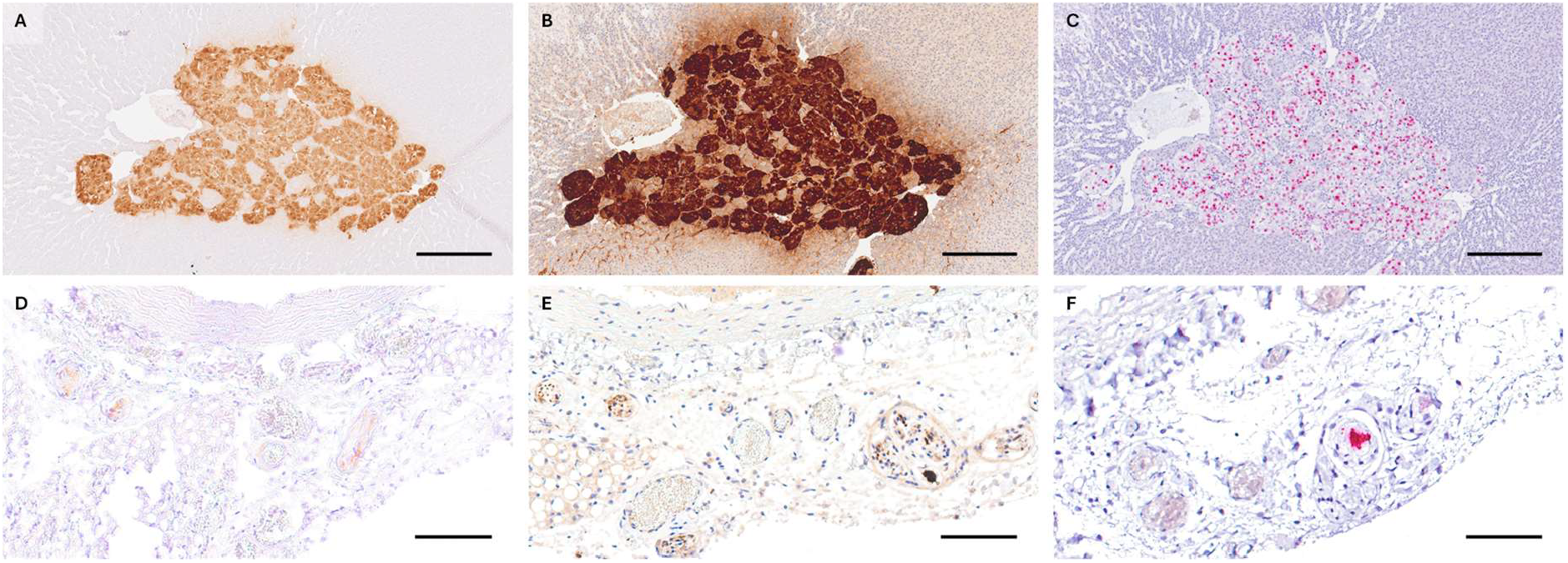
Comparison of three diagnostic methods, in the adrenal glands (A,B,C) and mediastinal nerve branches (D,E,F) of case 5. Diffusely positive chromaffin cells and multifocal positive nerve branches respectively, are seen in IHC using anti-BoDV-1 N mouse monoclonal antibody Bo18 (A and D), rabbit anti-BoDV-1 N polyclonal hyperimmune serum #201 (B and E) and RNAscope® ISH (C and F). Scale bars A-C: 250 μm and D-F:100 μm. Stain, A,B,D,E: DAB, hematoxylin counterstain. C,F: 2.5 HD Assay – RED, hematoxylin counterstain.

To confirm these findings, IHC using rabbit polyclonal hyperimmune serum #201 and RNAscope® ISH were performed for selected peripheral organs of case 5. While the staining patterns were comparable for the tested peripheral nerves and the adrenal medulla (Figure 7), the IHC Bo18 signal in the tubular epithelial cells of case 5 could not be reproduced by either of the confirmatory methods (data not shown).

### Phylogeographic analysis of BoDV-1 sequences from hedgehogs

Partial genomic sequences covering at least the BoDV-1 N, X and P genes (1,824 nt) were determined for all seven BoDV-1-positive hedgehogs. Phylogenetic analysis together with 258 BoDV-1 sequences derived from public databases (*1, 9*) revealed all seven sequences to belong to the BoDV-1 sequence subcluster 1A, which is in agreement with the origin of the animals from the southeast of Bavaria (Figure 8A, C). A more detailed analysis of subcluster 1A identified the hedgehog-derived sequences as belonging to subclades 1A.SE-1 (cases 1, 3, 5; all from district Rottal-Inn), 1A.SE-2 (case 7; Rottal-Inn) and 1A.SE-3 (cases 2, 4, 6; all from district Ebersberg; Figure 8B, D). In all cases, sequences of the same subclade derived from infected shrews, domestic mammals or humans were found in the same or neighboring districts as the respective hedgehog cases (Figure 8D).

**Figure 8.**
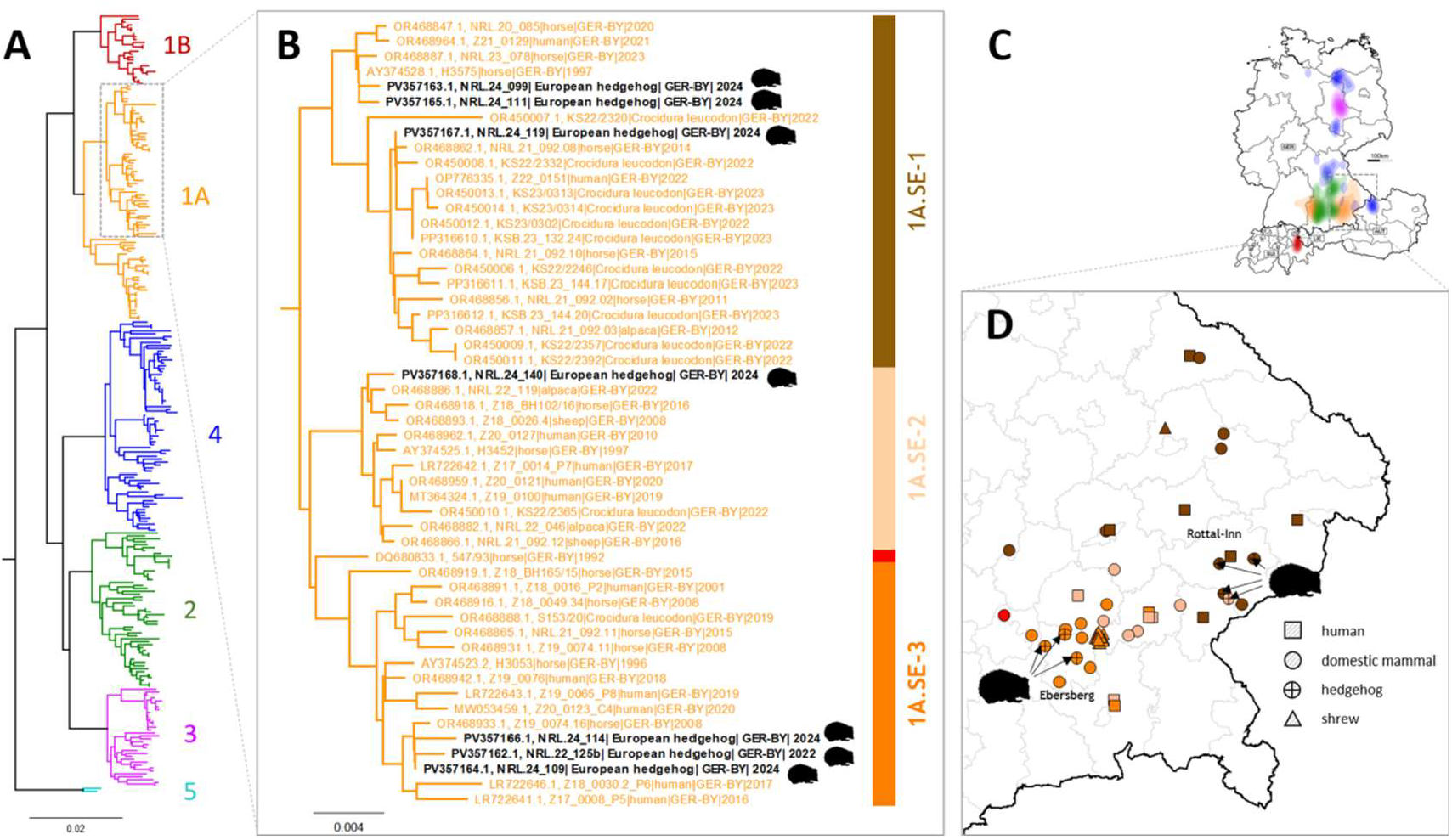
Phylogeographic analysis of BoDV-1 infections in hedgehogs in southeast Bavaria. (A) Phylogenetic analysis of partial genomic BoDV-1 sequences (N, X and P gene; 1,824 nucleotides, representing genome positions 54 to 1,877) of all seven BoDV-1-infected hedgehogs in combination with 258 BoDV-1 sequences from naturally infected animals and humans available from public databases (*1, 9*). Colors of tree branches represent BoDV-1 sequence clusters. (B) Detailed presentation of the subtree of cluster 1A containing all seven hedgehog-derived sequences (depicted in black). Colors of vertical bars represent subclades of cluster 1A as defined by Ebinger et al. (*1*) (C) BoDV-1-endemic area in Germany, Austria, Switzerland and Liechtenstein as determined by Ebinger et al. (*1*). Colors represent the phylogenetic clusters shown in panel A. (D) Detailed phylogeographic mapping of BoDV-1 cases from the phylogenetic subtree of cluster A shown in panel B to south-eastern Bavaria. Colors represent the different subclades. For data protection, human cases are mapped no more precise than to the center of the administrative district of their origin.

## Discussion

This report documents the first evidence of BoDV-1 infection causing meningoencephalomyelitis, radiculitis and in one case also focally extensive ganglioneuritis in wild European hedgehogs. While BoDV-1 is known to cause fatal encephalitis in a broad range of mammalian species, well-documented cases in wild mammals are so far restricted to a single European beaver (*Castor fiber*) from 2013 (*28*). The first case of the series described here was detected in 2022 before six additional hedgehogs from the same, relatively restricted area within a BoDV-1-endemic region in Bavaria, southern Germany were submitted in 2024 within only four months. We cannot exclude that this temporal and regional accumulation may represent a local emergence or increase of BoDV-1 infections of hedgehogs. However, it appears more likely that the initial cases raised the awareness of disease surveillance centers, leading to more frequent diagnosis of a previously underreported entity.

Our findings suggest that hedgehogs may be particularly susceptible to BoDV-1 infection, possibly due to their biological relationship to shrews, the reservoir hosts of BoDV-1 (*20*). This raised the question of whether infected hedgehogs may serve only as spill-over dead-end hosts that develop disease without viral shedding, or whether they may also show broad viral tissue distribution, viral shedding or even an asymptomatic infection, similar to BoDV-1 reservoir hosts (*11*). Such an intermediate role has been described for psittacines affected by parrot bornaviruses (PaBV-1 to -8; species *Orthobornavirus alphapsittaciforme* and *Orthobornavirus betapsittaciforme*). Affected birds suffer from neurological disease and, at the same time, they can transmit the virus to a broad range of other Psittaciformes that are originating from different continents and are thus unlikely to all represent original reservoir hosts of these viruses (*29*).

So far, all BoDV-1-positive hedgehogs presented with neurological signs induced by a lymphoplasmohistiocytic meningoencephalitis, similar to that of spill-over hosts such as horses, alpacas, sheep and humans (*30–32*). As this may be biased by our initial case selection focusing on animals with neurologic abnormalities, we extended the study to non-encephalitic hedgehogs from endemic areas. Hitherto, we could not detect BoDV-1 in brains of 23 non-encephalitic hedgehogs. However, a larger survey of neurologically inconspicuous hedgehogs in BoDV-1-endemic areas is required to rule out the possibility of mild or asymptomatic BoDV-1 infections.

Notably, BoDV-1-infected hedgehogs described herein exhibit differences from other affected species in terms of lesion and virus distribution spatial characteristics within the CNS. In horses, the most prominently affected CNS region is the hippocampal formation, followed by limbic system, basal ganglia, and brainstem (*31, 32*). Humans seem to consistently show virus infestation hotspots in the brainstem plus telencephalon or diencephalon (*30*). No such signature was seen in the hedgehogs. Instead, they showed a uniform distribution of inflammatory infiltrates, virus antigen and RNA throughout the entire CNS. Beyond neuronal infection, both IHC and ISH revealed a prominent infection of glial cells and Schwann cells, while viral RNA was also detected in ependymal cells of four animals (Figure 1). A similar cell tropism was previously described in the CNS of humans (*30*) and experimentally infected rats (*33, 34*). In the studied hedgehogs, inflammation was mainly restricted to the CNS and spinal nerve roots, while only one animal showed ganglioneuritis of the cranial mesenteric ganglion To our knowledge, distal ganglioneuritis is not a typical feature in naturally BoDV-1-infected horses or alpacas (*8, 31*), but it is a hallmark of PaBV infection in parrots (*29*). Inflammation in peripheral nerves has also been described for few BoDV-1-infected human patients, who presented with Guillain Barré-like neuropathy at early stages of infection (*5, 30, 35*).

Detection of BoDV-1 antigen or RNA in peripheral nerves and ganglia has been sporadically described also for alpacas and human patients as well as the BoDV-1-infected beaver (*28, 30, 31*). It has been discussed mainly as representing centrifugal virus dissemination from the brain, as experimentally shown for mice and rats (*36, 37*). Compared to these species, BoDV-1-positive cells in the PNS were surprisingly common in the analyzed hedgehogs, with positive nerve fibers and ganglion cells identified in several organ for each of them.

In the two animals with the most widespread viral tissue distribution based on RT-qPCR results, viral antigen was also seen in chromaffin cells of the adrenal medulla, while in one of them, focal cytoplasmic immunopositivity was even observed in a focus of renal tubular epithelial cells. Although the latter finding could not be confirmed by IHC using another anti-BoDV-1 N antibody or by RNAscope® ISH, it raises the concern that individual BoDV-1-infected hedgehogs might actually shed the virus via mucosal surfaces.

Unfortunately, urine, feces and mucosal swabs were not available from this animal. However, it has to be emphasized that the extent of viral presence on the epithelial surfaces of this single animal is much lower as compared to infected bicolored white-toothed shrews, in which viral antigen is usually found widespread on various epithelial surfaces (*11*).

So far, all BoDV-1-infected hedgehogs showed neurological signs before capture and there is no indication that they got infected by or infected other hedgehogs housed in the same rescue centers. In line with this observation, the BoDV-1 sequences found in the seven cases belonged to three different phylogenetic subclades, reflecting the dominant virus type in the region in which the animal was found (*1*). This argues for individual spill-over events from the local shrew reservoir, rather than for a hedgehog-adapted BoDV-1 variant circulating in their populations.

In summary, we present a series of Borna disease cases in European hedgehogs. This suggests that BoDV-1 infections may be significantly underreported in this species as well as wild mammals in general. It is therefore essential to consider BoDV-1 infection a possible differential diagnosis in hedgehogs with CNS signs and encephalitic lesions in endemic regions, even if Joest-Degen inclusion bodies are not present. Despite being distant relatives of bicolored white-toothed shrews, the identified BoDV-1-infected hedgehogs showed the signature of typical spill-over dead-end hosts, with fatal lymphoplasmohistiocytic encephalitis and an almost exclusively neurotropic infection. However, the broader viral presence across the PNS of hedgehogs and occasional detection of viral antigen in non-neural cells, possibly including renal epithelial cells in one animal, raise concerns as to whether singular infected hedgehogs may be able to shed virus. In such cases, the amount of excreted virus would likely be considerably lower as compared to regular reservoir hosts. Moreover, BoDV-1 spill-over transmission to humans appears to be generally inefficient, with only few cases per year even in areas in which the virus is broadly present in the local shrew population (*1, 9*). However, given the potentially close contact of humans and hedgehogs and the high case-fatality rate of zoonotic BoDV-1 infections, our results not only call for further investigations into the epidemiology of BoDV-1 infections in hedgehogs but also emphasize that standard hygiene measures should be implemented whenever handling hedgehogs and particularly for those with neurologic disorders.

## Supporting information

Supplemental table S1, Supplemental table S2,Supplemental table S3, appendix section B, appendix section C

## Acknowledgements

The authors would like to thank Sandra Aumiller, Sabine Brenner, Friedrich Faßler, Natalia Gerling, Gudrun Goldmann, Johann Gschlößl, Maximilian Hanslik, Weda Hoffmann, Moritz Maier, Astrid Nagel, Marina Schneider, Marion Segl, Juliane Stephan, Kathrin Steffen, Philip Starcky and Karin Stingl for their excellent technical assistance.

## Disclaimers

Part of the study was funded by the Bavarian State Ministry of Health, Care and Prevention (StMGPP), project Zoonotic Bornavirus Focal Point (ZooBoFo) Bavaria.

