## Supplemental table S1, Supplemental table S2,Supplemental table S3, appendix section B, appendix section C for "Borna disease virus 1 infection causing fatal meningoencephalomyelitis in wild European hedgehogs in known endemic areas, Germany, 2022 to 2024"

A. **Supplemental tables**

**Supplemental table S1. European hedgehogs tested for BoDV-1 in this study**

| Lymphohistiocytic encephalitis | Origin from BoDV-1-endemic area | Years | Number of tested animals | Number of BoDV-1 RT-qPCR-positive animals (%) |
| --- | --- | --- | --- | --- |
| Yes | Yes | 2022-2024 | 8 | 7 (88%) |
| No | Yes | 2024 | 23 | 0 |

**Supplemental table S2. Primers and probes used for RT-qPCR testing and sequencing of partial BoDV-1 genomes.**

| **Assay** | **Primer/Probe name** | **Sequence (5’ to 3’)** | **Reference** |
| --- | --- | --- | --- |
| BoDV-1 Mix-1 | BoDV-1_1258+ | TAGTYAGGAGGCTCAATGGCA | Schlottau *et al.*(*1*) |
|  | BoDV-1_1316_FAM | FAM-AAGAAGATCCCCAGACACTACGACG-BHQ1 | Schlottau *et al.*(*1*) |
|  | BoDV-1_1419- | GTCCYTCAGGAGCTGGTC | Schlottau *et al.*(*1*) |
| BoDV-1 Mix-6 | BoDV-1_2231+ | CAATYAATGCAGCYTTCAATGTCTT | Schlottau *et al.*(*1*) |
|  | BoDV-1_2285as_FAM | FAM-CCARCACCAATGTTCCGAAGCCG-BHQ1 | Schlottau *et al.*(*1*) |
|  | BoDV-1_2305- | GAATGTCYGGGCCGAGAG | Schlottau *et al.*(*1*) |
| panRusV-2a | RusV_234+ | CCCCGTGTTCCTAGGCAC | Thilen *et al.* (*2*) |
|  | RusV_256_P | FAM-GTGAGCGACCACCCAGCACTCCA-BHQ1 | Thilen *et al.* (*2*) |
|  | RusV_323- | TCGCCCCATTCWACCCAATT | Thilen *et al.* (*2*) |
| TBE1-Mix | TBEV_11054+ | GGGCGGTTCTTGTTCTCC | Schwaiger *et al.*(*3*) |
|  | TBEV_11073_P | FAM-TGAGCCACCATCACCCAGACACA-BHQ1 | Schwaiger *et al.* (*3*) |
|  | TBEV_11121- | ACACATCACCTCCTTGTCAGACT | Schwaiger *et al.* (*3*) |
| beta actin mix 2 * | ACT_F_1005-1029 | CAGCACAATGAAGATCAAGATCATC | Toussaint *et al.* (*4*) |
|  | ACT_P_1081-1105_HEX | HEX-TCGCTGTCCACCTTCCAGCAGATGT-BHQ1 | Toussaint *et al.* (*4*) |
|  | ACT_R_1135-1114 | CGGACTCATCGTACTCCTGCTT | Toussaint *et al.* (*4*) |
| eGFP mix 1 * | EGFP-1-F | GACCACTACCAGCAGAACAC | Hoffmann *et al*.(*5*) |
|  | EGFP-Probe1_HEX | HEX-AGCACCCAGTCCGCCCTGAGCA-BHQ1 | Hoffmann *et al.*(*5*) |
|  | EGFP-2-R | GAACTCCAGCAGGACCATG | Hoffmann *et al.*(*5*) |
| BoDV-1 amplification for sequencing | PaBV-2_1+ | TGTTGCGGTAACAACCAAC | Rubbenstroth *et al.* (*6*) |
|  | BoDV-1_1161- | TTAGACCAGTCACACCTATC | Schulze *et al.* (*7*) |
|  | BoDV-1_1068+ | GTATAGGCGCCGCGAGATAT | Schulze *et al.* (*7*) |
|  | BoDV-1_2311- | AAGATCGAATGTCTGGGCCG | Schulze *et al.* (*7*) |
| sequencing primers | BoDV-1_523+ | GCAGGAGCCGARCAGATCAAG | Schulze *et al.* (*7*) |
|  | BoDV-1_656- | GGTTGGCCGTTAATCCAATC | Schulze *et al.* (*7*) |
|  | BoDV-1_1621+ | GAAACCATCCAGACAGCTCAG | Schulze *et al.* (*7*) |
|  | BoDV-1_1816- | GAGGTGCAGGATGGGAGGG | Schulze *et al.* (*7*) |

*) Control reactions for the detection of host RNA (beta actin) or an external RNA spiked into the sample during RNA extraction (eGFP).

**Supplemental table S3. Additional information on BoDV-1-infected European hedgehogs included in this study.**

|  | Case 1 | Case 2 | Case 3 | Case 4 | Case 5 | Case 6 | Case 7 |
| --- | --- | --- | --- | --- | --- | --- | --- |
| Weight (g) | 960 | 800 | 830 | 640 | 480 | 542 | 784 |
| Sex | Male | Male | Male | Male | Female | Female | Male |
| Mode of death | Euthanized | Deceased | Euthanized | Euthanized | Euthanized | Euthanized | Euthanized |
| Neurological signs |  |  |  |  |  |  |  |
| Incoordination | x |  | x | x | x | x | x |
| Gait abnormalities | x | x |  | x |  | x | x |
| Seizures |  | x |  |  |  |  |  |
| Head tilt |  |  |  | x |  |  |  |
| Twitching |  | x |  |  | x |  |  |
| Apathy |  | x |  | x |  | x |  |
| Impaired thermo-regulation |  |  |  | x |  | x |  |
| Therapy |  |  |  |  |  |  |  |
| Treatment attempts | Insecticides, anthel-minthics, antibiotics, single dose of cortisone | No treatment | Not specified | Unspecified symptom-matic treatment | Not specified | Anthel-minthics and unspecified symptom-matic treatment | Not specified |

**B. Detailed description of immunohistochemical procedures**

The detailed immunohistochemical procedures for all used antibodies are described below. All antibodies had been tested and optimized for the use in this species in a pilot trial (data not shown). Immunohistochemistry for GFAP, CD3, Iba1, BoDV-1 (monoclonal antibody Bo18), rabies virus, canine distemper virus (CDV) was performed manually, while for Pax5 and BoDV-1 (rabbit polyclonal hyperimmune serum #201) an automated immunostainer was employed.

**B.1. Manual immunohistochemistry:**

**I. Deparaffinization and rehydration**

Sections were deparaffinized in xylene and rehydrated in descending ethanol series followed by rinsing with distilled water or Tris-buffered saline (TBS).

**II. Antigen retrieval**

Antigen retrieval was performed through heat-induced epitope retrieval (HIER) using a microwave, with an addition of Tris-EDTA buffer (pH 9.0) for CD3, an addition of citrate buffer (pH 6.0) for GFAP, Iba1 and CDV, and without pre-treatment for Bo18, while no antigen retrieval was performed for rabies virus. After the microwave, the sections were let to cool and then rinsed with distilled water or TBS.

**III. Blocking Endogenous Peroxidase Activity**

Endogenous peroxidase activity was quenched by incubating sections in 3% hydrogen peroxide (H_2_O_2_), followed by rinsing with TBS.

**IV. Blocking Non-Specific Binding**

Non-specific binding was blocked using ImPRESS Normal Horse Serum for GFAP, a stock solution comprising blocking buffer with goat serum (1:40 dilution) supplemented with Avidin for CD3 and CDV, a stock solution comprising blocking buffer with rabbit serum (1:40 dilution) supplemented with Avidin for Iba1, normal goat serum (1:20 dilution) for Bo18 and rabies virus.

Blocking buffer refers to a solution of TBS prepared by diluting a 10X TBS stock 1:10 with distilled water to obtain a 1X TBS working solution, with addition of Bovine Serum Albumin (BSA) (1%), Triton X-100 (0.1%), of Goldfish Gelatin (0.2%) and of a 10% solution of Sodium Azide (0.02%).

**V. Primary Antibody Incubation**

Sections were incubated with the primary antibodies as follows:

1. Rabbit polyclonal anti-GFAP antibody (Dako Denmark A/S, Denmark) in 1:800 dilution overnight at 4°C, diluted in blocking buffer.
2. Rabbit polyclonal anti-CD3 antibody (Dako Denmark A/S, Denmark) in 1:100 dilution for 1 hour at room temperature, diluted in stock solution (blocking buffer and goat serum in 1:40 dilution) supplemented with Biotin.
3. Mouse monoclonal Bo18 antibody (1:1,000 dilution) for 1 hour at room temperature, diluted in TBS.
4. Rabbit polyclonal anti-Iba1 antibody (Abcam, USA) in 1:500 dilution overnight at 4°C, diluted in blocking buffer supplemented with Biotin.
5. Mouse monoclonal anti-CDV antibody (VMRD, Pullman, WA, USA) in 1:1,000 dilution overnight at 4°C, diluted in stock solution (blocking buffer and goat serum in 1:40 dilution) supplemented with Biotin.
6. Mouse monoclonal anti-rabies virus antibody at a 1:200 dilution overnight at 4°C, diluted in TBS.

All sections were subsequently rinsed with TBS.

**VI. Secondary Antibody Incubation**

Sections were incubated with the secondary antibodies as follows:

1. GFAP: Sections were incubated with ImPRESS HRP anti-rabbit Polymer Kit for 30 minutes.
2. CD3: Sections were incubated with biotinylated goat anti-rabbit secondary antibody (1:200 dilution) for 50 minutes.
3. Bo18: Sections were incubated with biotinylated goat anti-mouse secondary antibody (1:200 dilution) for 50 minutes.
4. Iba1: Sections were incubated with biotinylated goat anti-rabbit secondary antibody (1:200 dilution) for 1 hour.
5. CDV: Sections were incubated with biotinylated goat anti-mouse secondary antibody (1:200 dilution) for 50 minutes.
6. Rabies: Sections were incubated with biotinylated goat anti-mouse secondary antibody (1:200 dilution) for 1 hour.

All sections were subsequently rinsed in TBS.

**VII. ABC Complex**

All sections except for GFAP were incubated with ABC reagent (Avidin-Biotin Complex), in a 1:100 dilution for 30 minutes, and subsequently rinsed in TBS.

**VIII. Detection, Counterstaining, Dehydration and Mounting**

Chromogenic detection was performed using 3,3'-diaminobenzidine (DAB) substrate. Counterstaining was performed using hematoxylin and sections were then rinsed in tap water. Afterwards dehydration was performed with an ascending ethanol series, sections were then cleared in xylene, and coverslipped using a permanent mounting medium.

**IX. Positive controls**

Sections of dog spinal cord (GFAP), dog lymph node (CD3 and Iba1), BoDV-1-infected horse brain (Bo18), rabies virus-infected dog brain (rabies Virus) and CDV-infected dog brain (CDV) were used as positive controls for the staining.

**B.2. Automated immunohistochemistry:**

Immunohistochemistry for Pax5 and BoDV-1 rabbit polyclonal hyperimmune serum #201 was performed using a fully automated staining system (BOND-III; Leica Biosystems, Wetzlar, Germany). Antigen retrieval was performed through heat-induced epitope retrieval (HIER), for Pax5 in a temperature of 60°C for 60 minutes, with an addition of citrate buffer (pH 6.0), and for BoDV-1 in EDTA buffer at 100°C for 100 minutes. The slides were incubated with primary antibodies anti-Pax5 (monoclonal, rabbit, dilution 1:50; Cell Marque, USA) and anti-BoDV1 (polyclonal, rabbit, dilution 1:5000) for 20 minutes and 32 minutes respectively. Antibody detection and counterstaining was performed through the BOND Polymer Refine Detection Kit (Leica Biosystems, Germany). Positive controls of human tonsils and human BoDV-1 infected brain, respectively, were used to confirm antibody specificity.

**C. Concurrent pathologies**

Five of the seven BoDV-1-positive hedgehogs histologically showed concurrent inflammatory changes of the lungs, ranging from mild lymphoplasmacytic interstitial pneumonia to lymphoplasmacytic and histiocytic peribronchitis and bronchopneumonia associated with intralesional nematodes, morphologically most consistent with *Capillaria aerophila* (*8*). Moreover, cases 1, 2, 3 and 7 showed a mild to moderate, multifocal, lymphoplasmacytic interstitial nephritis of unclear origin. In all cases, in which spleen was sampled, there was prominent extramedullary hematopoiesis, a finding common to this species (*9*).
